# Sharp tuning of head direction by somatosensory fast-spiking interneurons

**DOI:** 10.1101/2020.02.03.933143

**Authors:** Xiaoyang Long, Calvin K. Young, Sheng-Jia Zhang

## Abstract

Head direction (HD) information is intricately linked to spatial navigation and cognition. We recently reported the co-existence of all currently recognized spatial cell types can be found in the hindlimb primary somatosensory cortex (S1HL). In this study, we carried out an in-depth characterization of HD cells in S1HL. We show fast-spiking (FS), putative inhibitory neurons are over-represented in and sharply tuned to HD compared to regular-spiking (RS), putative excitatory neurons. These FS HD cells are non-conjunctive, rarely theta modulated, not locally connected and are enriched in layer 4/5a. Their co-existence with RS HD cells and angular head velocity (AHV) cells in a layer-specific fashion through the S1HL presents a previously unreported organization of spatial circuits. These findings challenge the notion that FS, putative inhibitory interneurons are weakly tuned to external stimuli in general and present a novel local network configuration not reported in other parts of the brain.

## Introduction

The ability to navigate from one place to another requires the knowledge of self in space. While hippocampal-entorhinal systems appear to map out the spatial environment (Hafting et al., 2005; O’Keefe and Dostrovsky, 1971; Solstad et al., 2008), the head direction (HD) system maintains an internally generated reference point to anchor our orientation in space. Cells that are tuned to HD were first reported in the rat postsubiculum (Taube et al., 1990a), which are strongly modulated by salient visual cues (Taube et al., 1990a, b). The neural substrate of HD cell activity has been shown to be dependent on vestibular input (Stackman et al., 2002; Stackman and Taube, 1997). The existence of cells responsive to the speed of angular head displacement (angular head velocity; AHV cells) and connection patterns between the lateral mammillary nucleus and the dorsal tegmental nucleus prompted the suggestion that a ring attractor network involving these structures may generate and propagate the HD signal throughout the brain (Clark and Taube, 2012).

The importance of the HD system in the spatial representation of the brain is highlighted by the persistence of HD representation upon manipulations that abolish position representation in hippocampal place cells (Bolding et al., 2019) and entorhinal grid cells (Brandon et al., 2011; Koenig et al., 2011). Particularly HD representations survive in cells that code HD conjunctively with grid or border representations (Brandon et al., 2011). Conversely, the elimination of HD signal into parahippocampal cortices through anteriodorsal thalamic nucleus lesions significantly disrupts HD, place (Calton et al., 2003; Harland et al., 2017; Stackman et al., 2002) and grid (Winter et al., 2015) cell signals. Therefore, available evidence suggests HD signaling goes beyond egocentric spatial representation and is also crucial for maintaining allocentric spatial representation.

It is known that if animals are deprived of sensorimotor input through passive transport by a cart, HD cells cannot maintain stable tuning (Stackman et al., 2003). If rotated passively, HD tuning may be intact but with greatly reduced firing rates (Knierim et al., 1998; Taube, 1995; Taube et al., 1990b) attributable to a mismatch of sensorimotor signals from insufficient head restraint (Shinder and Taube, 2011), suggesting self-motion is crucial for normal HD representation (Shinder and Taube, 2014). Similar properties exist for AHV cells, where passive rotation abolishes AHV in a subset of cells (Sharp et al., 2001). Collectively, these data point to the importance of sensorimotor input in the maintenance and persistence of HD activity.

Spatial selectivity in the brain has long been assumed to be the properties of specialized regions centered around the temporal cortex. More recent investigations suggest spatial representation may be more widespread than previously thought (Hok et al., 2018; Jankowski and O’Mara, 2015; Long and Zhang, 2018). We have recently discovered all currently recognized spatial cell types can be found in the rat hindlimb region of the primary somatosensory cortex (S1HL), including HD cells (Long and Zhang, 2018). In the current study, we carried out an in-depth characterization of S1HL HD and AHV cells. Remarkably, HD representation is not exclusive to regular-spiking (RS) putative principal cells. In fact, fast-spiking (FS) putative interneurons predominate HD representation with sharper tuning and appear to be enriched in layer 4/5a of S1HL and are not locally connected. AHV was also better tuned in FS and outnumber their RS counterparts, demonstrating a split layer distribution non-overlapping with FS HD cells. Our findings challenge the prevailing view that inhibitory interneurons are only broadly tuned to input stimuli and present a novel network configuration of, and a function for, HD representation in the brain.

## Results

### Somatosensory regular-spiking and fast-spiking HD cells

Putative single-cell recordings (**Supplementary Fig. S1**) through the S1HL (**Fig. 1a**) were obtained from seven rats performing a pellet chasing task in an open field (1 m x 1 m). As described previously (Long and Zhang, 2018), HD tuned cells can be found in the S1HL (**Fig. 1b**). The HD selectivity is stable within sessions (**Supplementary Fig. S2**) and is not biased by location (**Supplementary Fig. S3**). Neurons with mean vector length exceeding the 99^th^ percentile of the population distribution were classified as HD cells (**Fig. 1c**). To examine the relative contribution of putative excitatory and inhibitory neurons, we classified recorded neurons based on their spike widths and firing rates (**Fig. 1d**). From these criteria, we report 120/2107 (5.7%) cells being HD cells. Among HD cells, 62/120 (52%) were classified as regular-spiking (RS), 36/120 (30%) were classified as fast-spiking (FS) and 22/120 (18%) were unclassified (**Supplementary Fig. S4**). Only 44% of the RS HD cells were “pure” HD cells with no detectable conjunctive representation while 100% of the FS HD cells coded for HD exclusively (**Supplementary Fig. S4**). We did not find a difference in how well conjunctive and non-conjunctive cells coded for HD (**Supplementary Fig. S5**); hence we included all classified HD cells (i.e. conjunctive and non-conjunctive) for subsequent analyses. Both populations showed uniform distribution of HD tuning (χ^2^-test; *P* = 0.82 for FS HD cells and *P* = 0.81 for RS HD cells; **Fig. 1e**) From reconstructed tetrode tracks, the spatial distribution of RS and unclassified HD cells appear to be uniformly distributed across layers 3 to 6, while FS HD cells are enriched between layer 4 and 5a, avoiding deeper layers altogether (**Fig. 1f**).

**Fig. 1:**
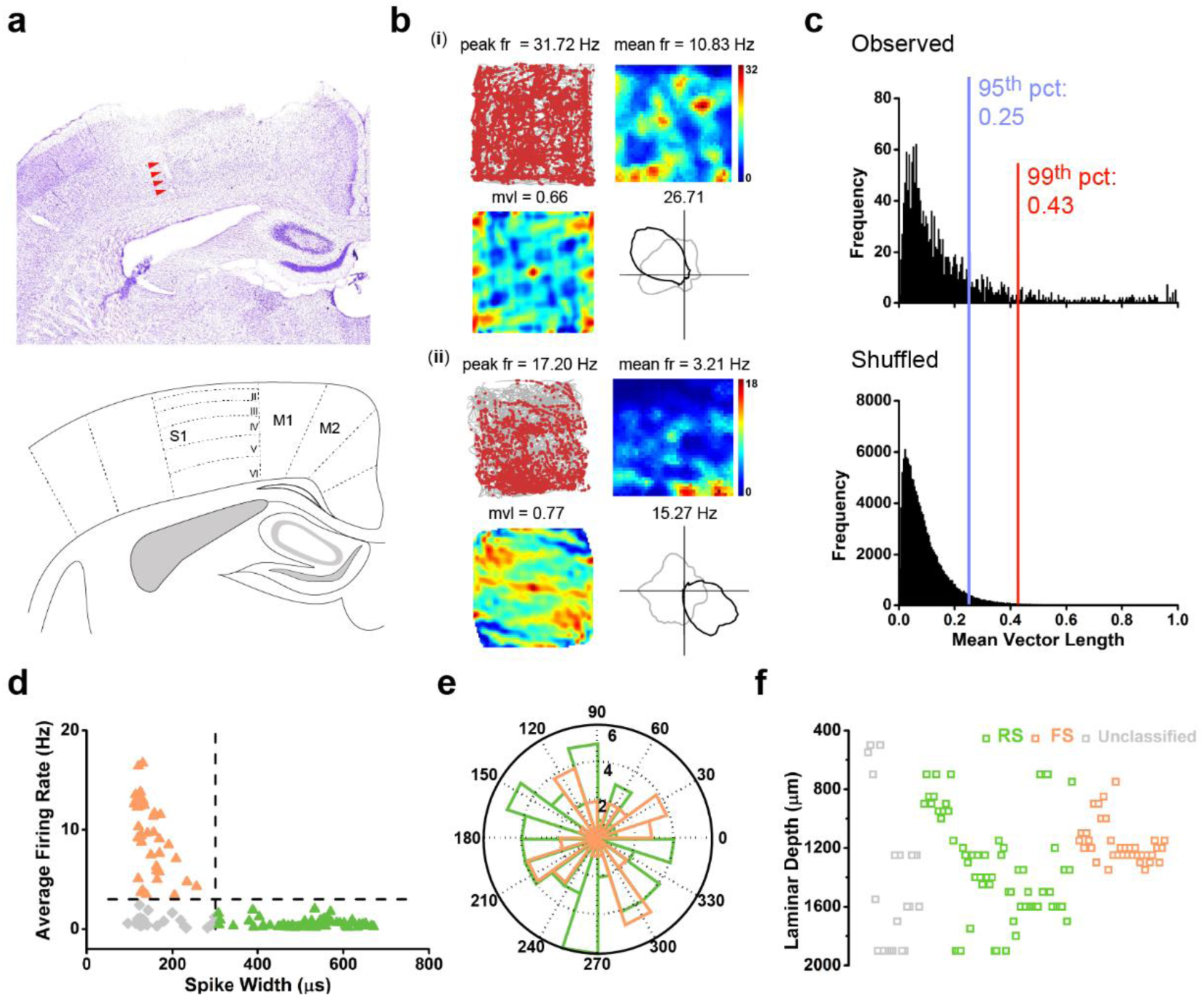
Both regular-spiking (RS) and fast-spiking (FS) cells are tuned to head direction in the S1HL. **a**, A representative Nissl-stained coronal section shows tetrode trajectory (indicated with red arrowheads) tracking through all of six layers across the rat primary somatosensory cortex (top). Idealized S1 layer boundaries are depicted on the reference atlas (bottom). **b**, Two representative somatosensory HD cells. Trajectory (grey line) with superimposed spike locations (red dots; top left), smoothed rate maps (top right), spatial autocorrelation (bottom left) and HD tuning curves (black) plotted against dwell-time (grey) in polar coordinates (bottom right). Firing rate is color-coded with dark blue indicating minimal firing rate and dark red indicating maximal firing rate.

The scale of the autocorrelation maps is twice that of the spatial firing rate maps. Peak firing rate (fr), mean fr, mean vector length (mvl) and angular peak rate for each representative HD cell are labeled at the top of the plots. **c**, Distribution of mean vector length for the entire pool of recorded somatosensory cells. The top panel shows the distribution for observed values. The bottom panel shows the distribution for randomly shuffled data from the same pool of recorded S1HL. **d**, Distribution of the average firing rate versus peak-to-through distance (spike widths) shows three classes of HD cells: regular-spiking (RS; green), fast-spiking (FS; orange) and the unclassified cells (gray). **e**, The circular distribution of the preferred direction for two discrete FS (orange) and RS (green) groups from all seven animals. **f**, Reconstructed recording depth for all S1HL HD cells.

### Sharply tuned fast-spiking HD cells

FS tuning to HD has been described as being weak within the canonical HD circuit (Peyrache et al., 2019; Tsanov et al., 2011; Tukker et al., 2015), and only a single study to date has reported FS HD tuning comparable to RS HD cells in the hippocampus (Leutgeb et al., 2000). Therefore, we sought to examine how HD tuning varied between RS and FS. Remarkably, we found that RS HD cells were generally less well-tuned (**Fig. 2a, c**) compared to FS HD cells (**Fig. 2b, d**). Given FS cells have higher firing rates which could potentially inflate HD tuning measures, we downsampled FS firing by randomly omitting spikes from the raw spike train to match the average firing rate of RS HD cells (**Fig. 2f**) to test if the sharper FS HD tuning persists (e.g. **Fig. 2e**). Downsampling did not change the mean vector length (**Fig. 2g**), tuning width (**Fig. 2h**) and angular stability (**Fig. 2i**) of FS HD cells and all these measures remained significantly higher than RS HD cells (**Fig. 2g-i**). These data collectively showed FS HD cells are better tuned than their RS counterparts, independent of spike number.

**Fig. 2:**
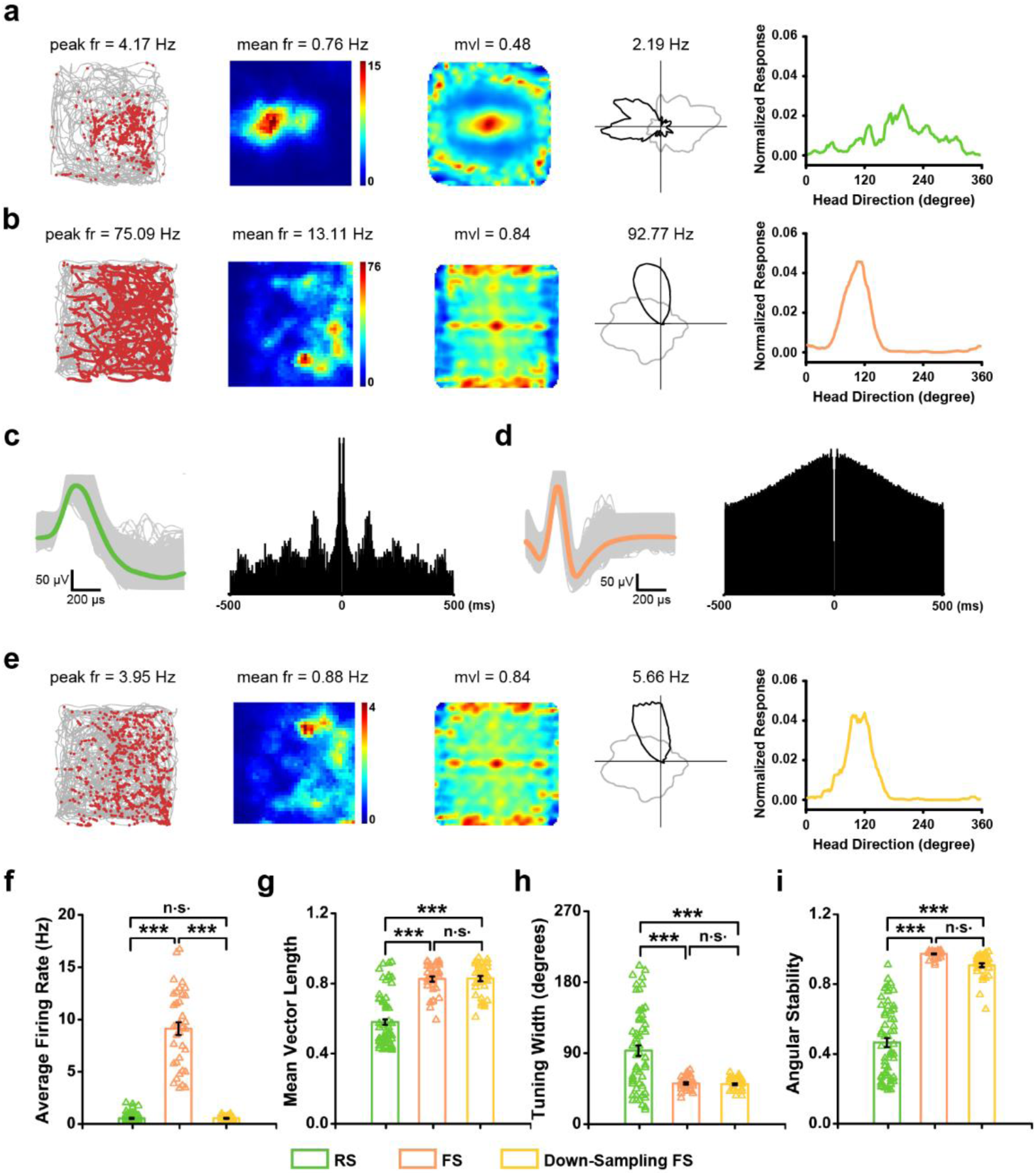
Sharper tuning of fast-spiking head direction cells. **a, b**, Two representative examples of somatosensory RS (**a**) and FS (**b**) HD cells. From left to right, trajectory (grey line) with superimposed spike locations (red dots) (left), smoothed rate maps, spatial autocorrelation, HD tuning curves (black) plotted against dwell-time and tuning curves are presented. Firing rate is color-coded with dark blue indicating minimal firing rate and dark red indicating maximal firing rate. The scale of the autocorrelation maps is twice that of the spatial firing rate maps. Peak firing rate (fr), mean fr, mean vector length (mvl) and angular peak rate for each representative head direction cell are labelled at the top of the panels. **c, d**, waveform of the representative RS HD cell (left in **c**) displays the longer spike duration than that of the representative FS HD cell (left in **d**). Representative 1000 ms spike-time autocorrelogram of the RS HD cell (right in **c**) shows a faster decay than that of the FS HD cell (right in **d**). **e**. Downsampled data for the FS HD cell depicted in (**b**). **f**, Comparable average firing rates between RS HD cells and downsampled FS HD cells. **g**, Mean vector lengths of FS HD cells (orange, *n* = 36) as well as downsampled FS HD cells (yellow, *n* = 36) are significantly higher than that of RS HD cells (green, *n* = 62). Mean vector lengths of FS HD cells do not differ between full-sampled and downsampled data. **h**, RS HD cells have wider tuning than FS HD cells and downsampled FS HD cells. **i**, FS HD cells and downsampled FS HD cells show higher angular stability than RS HD cells. Data are means ± s.e.m.; \*\*\**P* < 0.001, two-tailed unpaired *t*-test for RS HD vs FS HD cells or vs down-sampled FS HD cells; two-tailed paired *t*-test for FS HD cells versus down-sampled FS HD cells (**f**-**i**). green, RS; orange, FS; yellow, down-sampling FS.

### Weakly theta-modulated FS HD cells

Theta oscillations in the limbic system have been implicated in the integration of spatial inputs (Burgess and O’Keefe, 2011) and are necessary for the normal place (Bolding et al., 2019) and grid (Brandon et al., 2011; Koenig et al., 2011) cell activities. However, theta modulation of HD cells, in general, appears to be location- and cell type-dependent (Blair et al., 1999; Leutgeb et al., 2000; Preston-Ferrer et al., 2016; Sharp, 1996; Tsanov et al., 2011; Tukker et al., 2015). Therefore, we sought to characterize theta modulation of HD cells in the S1HL. Firstly, theta oscillations can be detected in the S1HL (**Fig. 3a**). An example of theta modulated RS HD cell autocorrelogram was shown in **Fig. 3c** and a non-theta modulated FS HD cell are shown in **Fig. 3d**. In general, spectral distribution of autocorrelograms of RS HD cells revealed a large peak at 7.5 Hz (**Fig. 3f**), while FS HD cell autocorrelograms had a nominal low frequency (< 2 Hz) peak (**Fig. 3g**). Most of the HD cells, RS or FS, had low percentage power in their spike autocorrelogram spectra (**Fig. 3e**). However, a subset of RS HD cells and a very few FS HD cells exhibited higher percentage theta power and were deemed to be theta modulated (**Fig. 3h**). In fact, there was a 15 fold difference in the proportion of RS (41.9%) and FS (2.8%) HD cells that were found to be theta modulated (**Fig. 3i**).

**Fig. 3:**
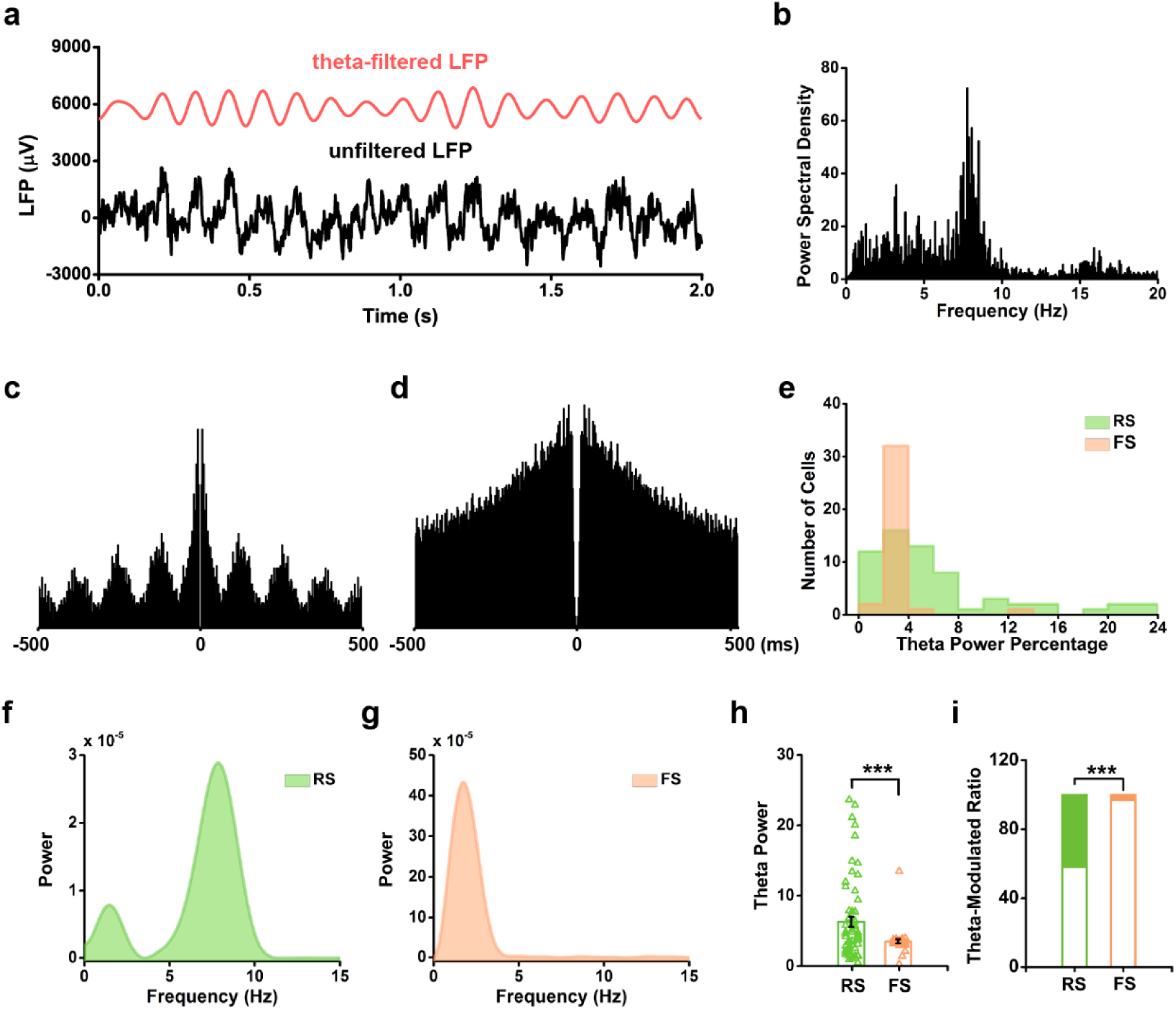
Fast-spiking head direction cells show little theta modulation. **a**, Prominent theta oscillations in the S1HL during locomotion. The unfiltered signal is in black and theta-filtered (4-11 Hz) signal is in red. **b**, Power spectral density for the whole recordings session shown in **a. c-d**, Two representative spike-time autocorrelograms of RS (**c**) and FS (**d**) HD cells. The RS HD cell shows clear theta modulation. **e**. Distribution of theta power percentage in the power spectrum of spike-time autocorrelograms of RS and FS HD cells. **f**-**g**. The power spectrum of spike-time autocorrelograms of the representative RS (**c**) and FS (**d**) HD cells, respectively. **h**. RS HD cells show significantly higher theta power percentage than FS HD cells. (mean ± s.e.m.; \*\*\**P* < 0.001, two-tailed unpaired *t*-test). **i**. The fraction of theta-modulated RS HD cells (26/62) was significantly higher than that of FS HD cells (1/36) (Z = -4.16, \*\*\**P* < 0.001, binomial test).

### FS HD cells in bursty discharge patterns

Bursting activity has been shown to play pivotal functions in spatial selectivity and orientation (Ebbesen et al., 2016). We found that FS HD cells exhibited bursting activity when the animals’ heads oriented in their preferred direction (**Fig. 4a**). The interspike intervals (ISI) histogram revealed the distinct temporal discharge patterns for RS and FS HD cells (**Fig. 4d**). Consistent with previous studies (Ebbesen et al., 2016; Latuske et al., 2015), the first two principal components of ISI distribution probability distinguished RS and FS cells (**Fig. 4b**). The bursty firing pattern of FS HD cells was also reflected in the cumulative probability distributions, with ISI being shorter than 20 ms (**Fig. 4c**).

**Fig. 4:**
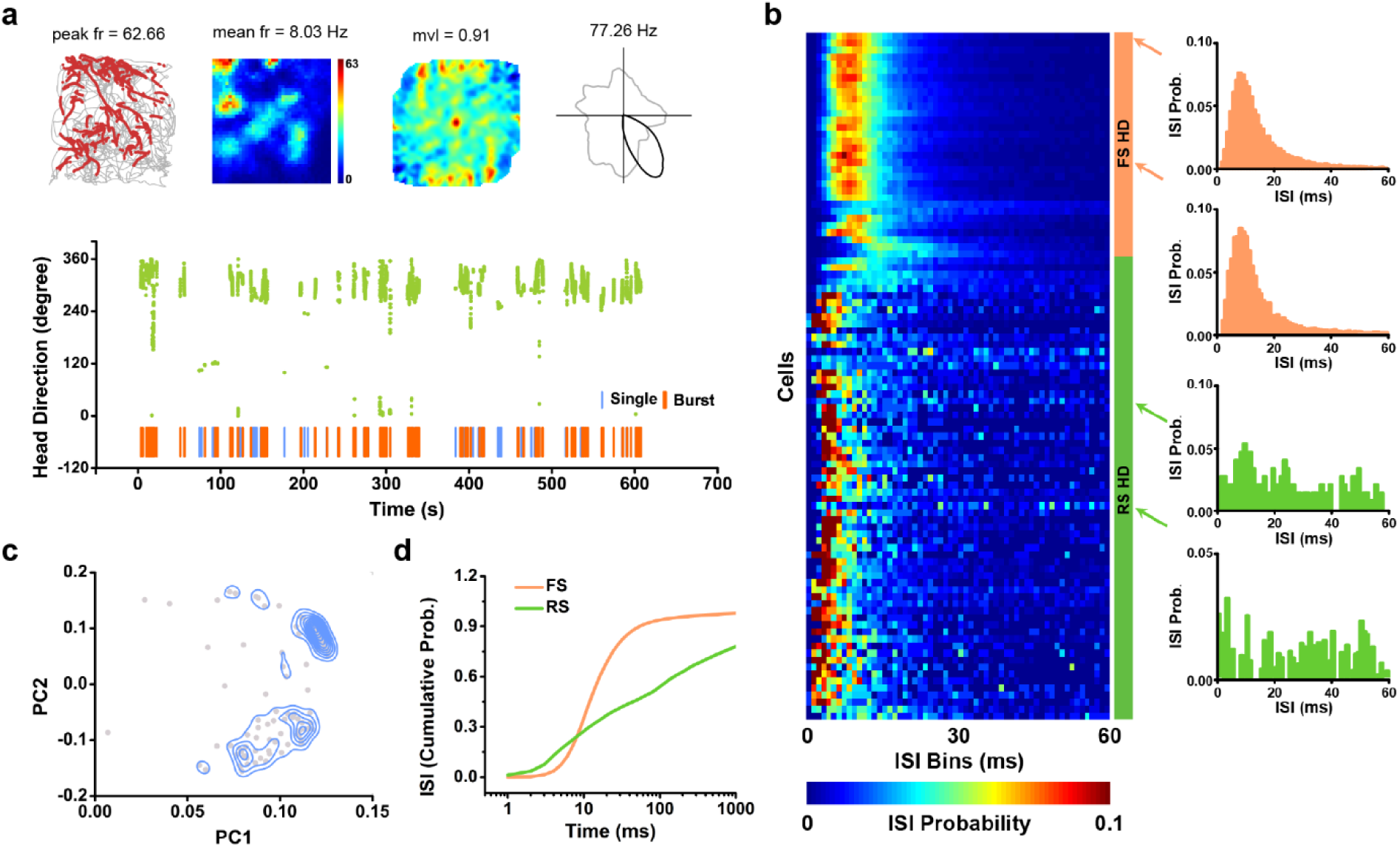
Fast-spiking head direction cells exhibit bursty firing mode. **a**, A representative example of FS HD cell discharging in a bursty pattern. From left to right, plots show: trajectory (grey line) with superimposed spike locations (red dots); smoothed rate maps; spatial autocorrelation; and head direction tuning (black) against dwell-time plot (grey). Firing rate is color-coded with dark blue indicating minimal firing rate and dark red indicating maximal firing rate. The scale of the autocorrelation maps is twice that of the spatial firing rate maps. Peak firing rate (fr), mean fr, mean vector length (mvl) and angular peak rate for each representative HD cell are labeled at the top of the panels. Below, the raster plot of the spike train emitted by the FS HD cell and corresponding HD (green dots). FS HD cell fires trains of burst spikes when the animal’s HD matches the cell’s preferred direction. Single spikes and bursty trains of spikes emitted by the FS HD cells are discriminated by ISI of 15 ms. **b**, ISI histograms (left) of pooled FS and RS HD cells, sorted by average firing rate. Representative ISI histograms of two somatosensory FS HD cells (upper right, orange) and two RS HD cells (lower right, green). FS HD cells tend to fire burst at 50-250 Hz (4-to 20-ms intervals). **c**, A principal component analysis is computed based on the ISI distribution. Scatterplot shows the first two principal components (PC1 and PC2 with 2D kernel smoothed density estimate in blue lines). **d**, Cumulative probability distribution of interspike intervals (ISI) for FS and RS HD cells. FS HD cells fired with ISI shorter than 20 ms.

### Somatosensory angular head velocity cells

The generation of HD selectivity is believed to involve AHV cells in proposed ring attractor models (Stratton et al., 2010; Zhang, 1996). A recent report showed that AHV cells can be found alongside HD cells in the neighboring motor cortex (Mehlman et al., 2019). Therefore, we hypothesized AHV cells should also be present in the S1HL. A total of 308/2107 putative single units recorded from the S1HL were classified as AHV cells (e.g. **Fig 5a, b**). Both symmetrical and asymmetrical AHV cells were found in the S1HL (**Fig. 5c, d**). As with HD cells, we further extended our AHV cell characterization with RS and FS AHV classification (**Fig. 5e**). Most AHV cells were non-conjunctive, and the most common conjunctive feature was HD (**Supplementary Fig. S6**), followed by location selectivity (i.e. exhibiting place cell-like selectivity; **Supplementary Fig. S7**). Almost half of the AHV cells were FS cells, which also had statistically significant higher AHV scores than RS AHV cells as a group (**Fig. 5f**). A large majority (85.7%) of RS AHV cells and a half (50%) of the FS AHV cells were found to be theta modulated (**Fig. 5h, j**); there appeared to be clear segregation of non-modulated and modulated cells based on our classification (**Fig. 5i**). Of note, a much larger proportion of FS AHV cells were found to be theta modulated than FS HD cells (**Fig. 5j**). Unlike the pattern seen for HD cells, RS AHV cells appear to dominate layers 3 while FS AHV cells were more common in layer 5 and deeper; layer 4 seemed to contain the least number of AHV cells (**Fig. 5g**).

**Fig. 5:**
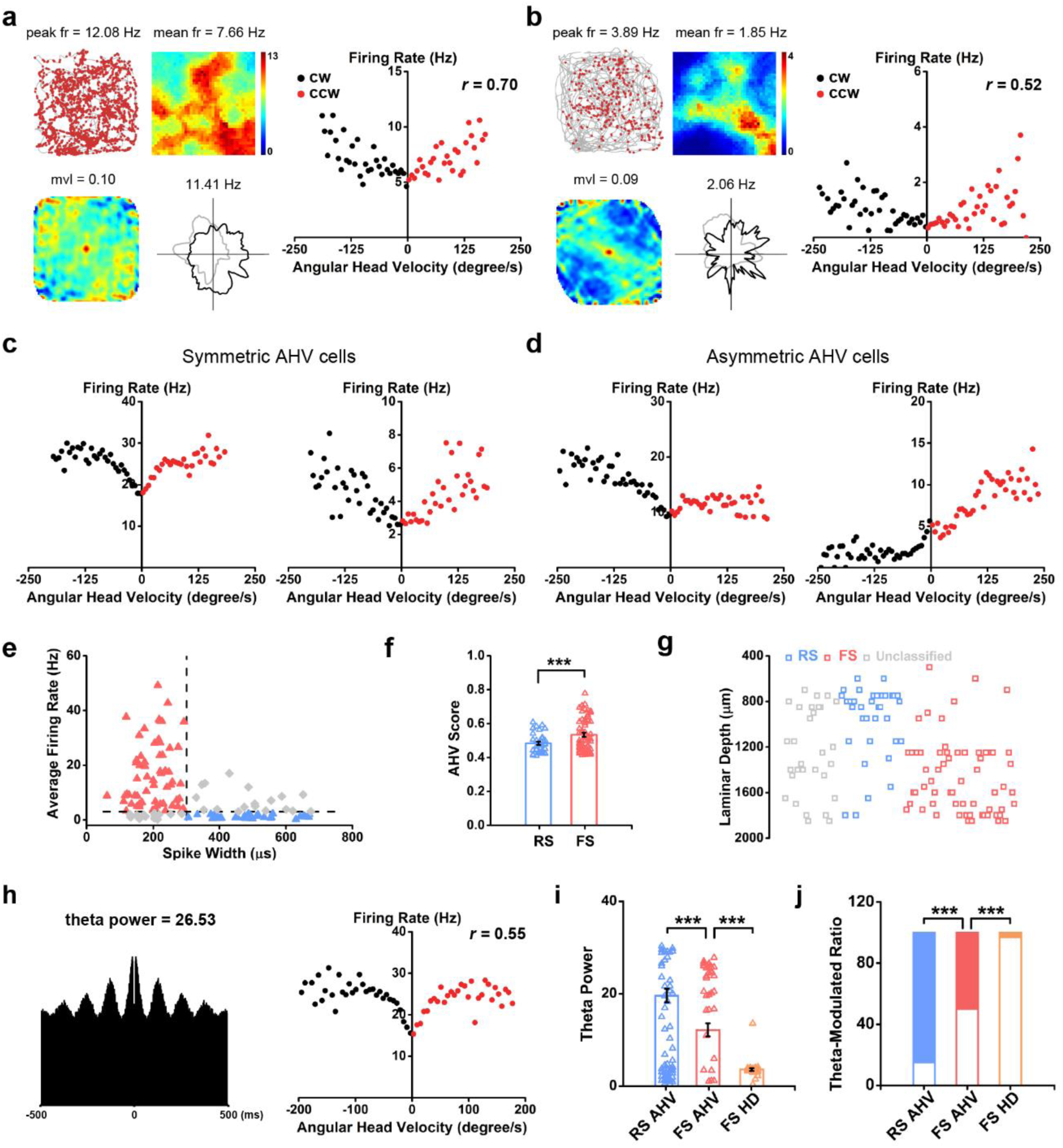
Somatosensory regular-spiking (RS) and fast-spiking (FS) angular head velocity cells. **a**, A representative FS AHV cell. Trajectory (grey line) with superimposed spike locations (red dots) (top left), smoothed rate maps (top right), spatial autocorrelation bottom left and HD tuning curves (black) plotted against dwell-time (grey; bottom right). Firing rate is color-coded with dark blue indicating minimal firing rate and dark red indicating maximal firing rate. The scale of the autocorrelation maps is twice that of the spatial firing rate maps. Peak firing rate (fr), mean firing fr, mean vector length (mvl) and angular peak rate for each representative head direction cell are labelled at the top of the panels. Right, the scatterplot of binned firing rates versus angular head velocity. **b**, The same as **a** but for a representative RS AHV cell. **c**, Two examples of symmetric AHV cells. **d**, Two representative asymmetric AHV cells. **e**, Distribution of the average firing rate vs peak-to-through distance (spike widths) defines three classes of AHV cells: RS (blue), FS (red) and unclassified (gray). **f**, The averaged AHV score of FS AHV cells is substantially higher than that those of RS AHV cells. Data are means ± s.e.m.; \*\*\**P* < 0.001, two-tailed unpaired *t*-test. **g**, The reconstructed depth distribution of recorded AHV cells. **h**, Representative example of somatosensory theta-modulated FS AHV cell. Left, the spike-time autocorrelogram; Right, the scatterplot of the firing rate versus the angular velocity of the representative AHV cell. Theta power and correlation coefficient (r) are labelled at the top of the panel. **i**, Theta power of FS AHV cells (red) compared to RS AHV cells and FS HD cells. **j**, The ratio of theta-modulated FS AHV cells (31/62, 50%) is significantly higher than that of FS HD cells (1/36, 2.8%) but lower than that of RS AHV cells (30/35, 85.7%). Data are means ± s.e.m.; \*\*\**P* < 0.001, ANOVA test (**i**); binomial test (**j**).

### No detectable putative monosynaptic connections involving FS HD cells

Lastly, given the apparent layer-specific distribution of HD and AHV cells in a cell-type dependent manner, we sought to identify putative monosynaptic connections (Bartho et al., 2004; Csicsvari et al., 1998; Latuske et al., 2015) for a better understanding of functional links between HD, AHV and other cells in the S1HL. We found a small number of RS HD cells made local excitatory (and sometimes reciprocal) connections to both simultaneously recorded RS and FS cells (27/458 pairs; 5.9%; **Fig. 6a**). Surprisingly, there were no identified putative monosynaptic connections between FS HD cells from 251 simultaneously recorded cell pairs. The spike-time cross-correlogram of two simultaneously recorded FS HD cells on the same tetrode showed no detectable synaptic connection (e. g. **Supplementary Fig. S8**). Comparable to RS HD cells, 5.8% of simultaneously recorded cell pairs (19/323) involving RS AHV cells made excitatory connections with both RS and FS cells – mainly with other AHV cells (**Fig. 6b**). A large proportion of cell pairs (7.7%; 41/532) involving FS AHV cells were found to make inhibitory connections to RS and FS, including that are tuned to AHV themselves (**Fig. 6c**). We were also able to detect putative common input-driven of FS AHV cells (**Fig. 6c-ii**).

**Fig. 6:**
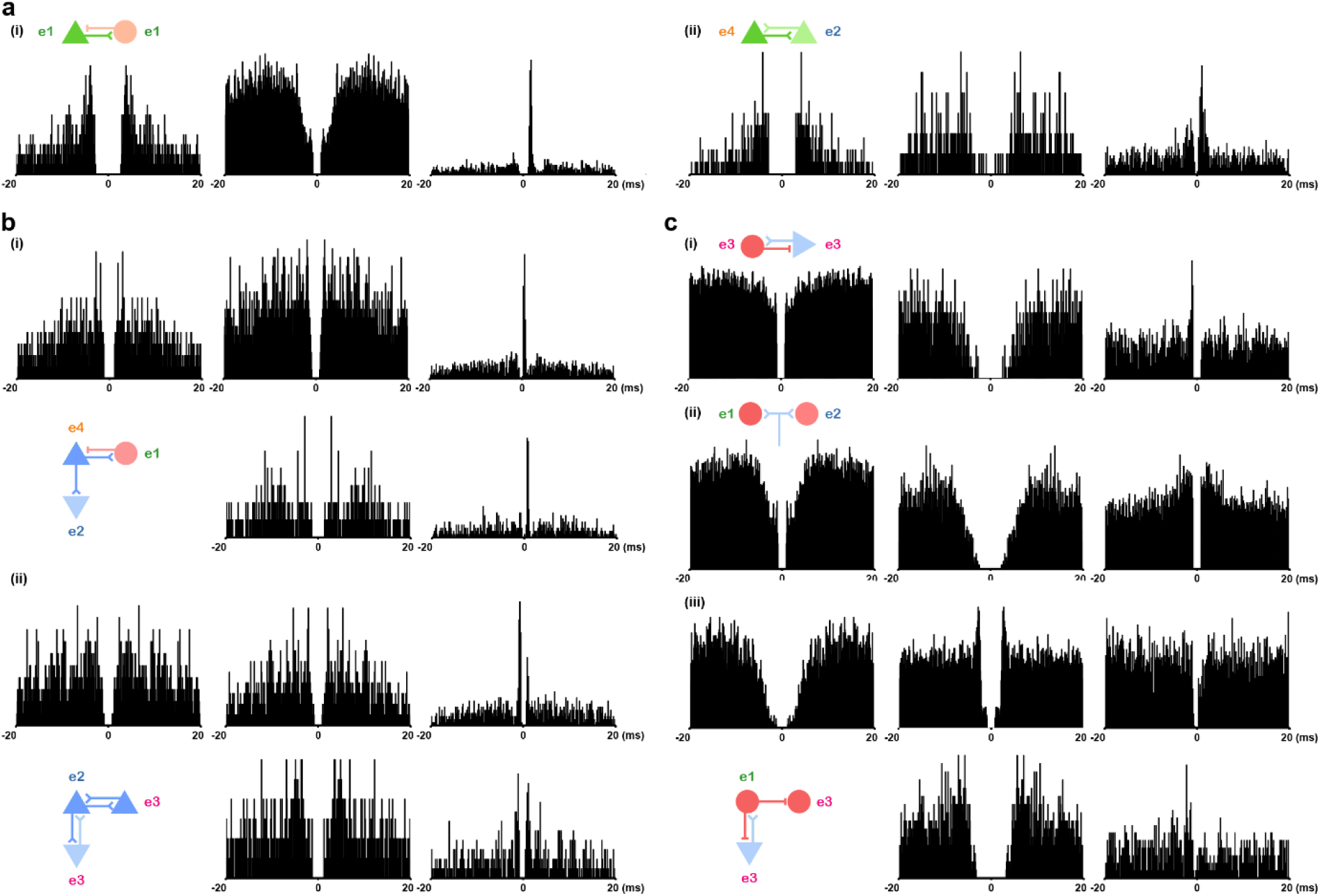
Putative monosynaptic connections between HD/AHV cells and simultaneously recorded neurons. **a**, Two representative monosynaptic connections as revealed by the spike-time cross-correlogram. Left, autocorrelogram of the reference cell; Middle, autocorrelogram of the target cell; Right, cross-correlogram. Hypothesized synaptic connectivity is indicated at the top left of each pair. The tetrode number (e1-e4) is labeled on the left corner of each representative cell. **b**, Putative monosynaptic connections between regular-spiking AHV cells and simultaneously recorded neurons. **c**, Putative monosynaptic connections between fast-spiking AHV cells and simultaneously recorded neurons.

## Discussion

Our previous report showed all currently recognized spatial cell types can be found in the S1HL, including HD cells (Long and Zhang, 2018). In this study, we include AHV as an additional spatial feature the S1HL cells also encode. In contrast to a predominant excitatory cell exhibiting HD tuning described elsewhere in the brain, we show disproportionate large numbers of putative interneurons code HD with higher precision than their putative excitatory neuron counterparts. Together, we present a novel configuration of HD circuitry in the S1HL that is dissimilar to those described previously in canonical HD systems.

### Fast-spiking cells are tuned to head direction in S1HL

While spatially related firing activities have been thought to be mostly confined to the hippocampal-entorhinal system (Moser et al., 2017), the existence of HD cells outside the canonical HD/spatial circuits is not new. Head direction selectivity has been reported in the lateral dorsal thalamic nucleus (Mizumori and Williams, 1993), the striatum (Mehlman et al., 2019; Mizumori et al., 2000; Ragozzino et al., 2001; Wiener, 1993), motor cortex (Mehlman et al., 2019; Mizumori et al., 2005), visual cortex (Chen et al., 1994) and the nucleus reuniens (Jankowski et al., 2014). Sharply tuned HD cells have been reported in the layer 2/3 of the MEC. These cells were not classified as FS or putative interneurons (Zutshi et al., 2018). The only reported putative interneuron tuning of HD is in the hippocampus, where HD cells appear to be exclusively FS cells (Leutgeb et al., 2000). Thus, our discovery of sharply tuned FS HD cells co-existing with less well-tuned RS HD cells represents a novel and unique observation specific to the S1HL.

### Fast-spiking cells are better tuned than regular-spiking cells to head direction

The most surprising finding in our study is that 10% of all recorded FS are HD cells, whereas only ∼5% of RS and unclassified cells are HD. In fact, if we restrict HD classification to include only non-conjunctive HD cells, FS HD cell proportionally outnumbers RS or unclassified HD cells 5:1 (10% vs 2%). Our data indicate RS HD cells are similar to those described elsewhere (Preston-Ferrer et al., 2016; Tsanov et al., 2011; Tukker et al., 2015), demonstrating: 1) firing characteristics of putative excitatory principal cells and; 2) are mostly theta-modulated. In past studies examining HD representation in the canonical HD circuit, FS cells were either excluded for spatial selectivity analysis (Coletta et al., 2018) or have been shown to weakly tuned to HD (Peyrache et al., 2019; Tukker et al., 2015) but better tuned to AHV (Preston-Ferrer et al., 2016). These past studies are consistent with previous work elsewhere in sensory and spatial systems, showing that FS cells are poorly tuned to manipulated input features (Atallah et al., 2012; Cardin et al., 2007; Grieves and Jeffery, 2017; Kato et al., 2013; Kerlin et al., 2010; Miyamichi et al., 2013; Zariwala et al., 2011) but may be better suited for coding speed (Gois and Tort, 2018; Kropff et al., 2015; Preston-Ferrer et al., 2016; Ye et al., 2018). However, in the cat visual cortex (Cardin et al., 2007; Runyan et al., 2010), mouse auditory cortex (Moore and Wehr, 2013), as well as in the motor cortex of monkeys (Merchant et al., 2008), FS/putative PV interneurons can exhibit stimulus selectivity comparable to principal cells. Particularly, Cardin and colleagues found all FS cells in layer 4, but not other layers, are sharply tuned to the spatial orientation of presented visual stimuli and are only marginally broader than RS cell tuning (Cardin et al., 2007). Given HD tuning across the whole population at any given HD is coherent (Peyrache et al., 2015), unlike in other systems where input selectivity is heterogeneous and overlapping (Isaacson and Scanziani, 2011), having divergent but homogeneous HD input may drive sharp FS HD tuning in the S1HL. However, our current data suggest FS HD cells are not locally connected, which indicates these FS cells may not be conventional PV+ neurons that have extensive local connections (Koelbl et al., 2015). High-density silicone probe recordings (Bartho et al., 2004; Csicsvari et al., 1998) and cell-type-specific tracing studies (Weible et al., 2010) may provide further insight on the identities and connectivity of FS HD cells reported here. To the best of our knowledge, no previous study has shown FS cells or putative PV+ interneurons displaying superior stimulus selectivity than their RS/putative excitatory cell counterparts. While the neurochemical identity of our FS HD cells remains to be elucidated, we provide the first *prima facie* evidence that putative PV+ interneurons can have sharply tuned feature selectivity.

### Generation of HD and AHV activity in the S1HL

The presence of AHV cells in the S1HL opens up the possibility that HD selectivity can be locally generated in S1HL. We show both symmetrical and asymmetrical AHV cells are more likely to be putative inhibitory interneurons (i.e. FS cells). Along with RS/ putative excitatory HD cells, all basic components of a theorized ring attractor are available within the S1HL for *de novo* HD signal generation (Stratton et al., 2010). However, none of the brain areas exhibiting HD tuning examine so far is independent of the canonical HD circuit (Bassett et al., 2007; Blair et al., 1998; Clark and Taube, 2012; Goodridge and Taube, 1997; Mehlman et al., 2019); our S1HL HD cells are unlikely to be different. In our previous report, we suggested that S1HL spatial selectivity is likely to be an efferent copy inherited from elsewhere, possibly from motor areas given their extensive functional and anatomical connections (Long and Zhang, 2018). HD and AHV signals from the motor cortex (Mehlman et al., 2019; Mizumori et al., 2005; Wiener, 1993) may arrive through projections to layer 5 (Veinante and Deschenes, 2003), consistent with our observation of FS HD cell enrichment at the border of layer 4/5. Layer 4 is a focal point for thalamocortical input via the ventral posterolateral nucleus (VPL). PV expression can be detected at highest levels across layers 4 and 5 in mice (Almasi et al., 2019; Meyer et al., 2011; Narayanan et al., 2017) and appear to be selectively enriched in layer 4 in rats (Bodor et al., 2005), where thalamic inputs strongly target and activate PV+ neurons (Cruikshank et al., 2007; Sermet et al., 2019) in the vibrissae S1. Thalamic afferents, particularly from VPL where vestibular inputs have been reported (Nagata, 1986), may constitute a novel alternative pathway for establishing HD signalling in the S1HL.

### Functional significance of sharply tuned FS HD cells in the S1HL

In the S1HL, the general rule of FS are only weakly or broadly tuned to sensory input holds true for tactile stimuli (Hayashi et al., 2018; Murray and Keller, 2011), but our data suggest HD representation is sharply tuned. What is the functional significance of having putative inhibitory interneurons sharply tuned to HD? Firstly, it is entirely possible that these sharply tuned putative interneurons carry out the same proposed function elsewhere in the brain – to further refine principal cell HD representation. Although we have shown that RS HD representation is largely less well-tuned than FS cells, it is entirely possible that RS HD representation may be weaker without FS HD refinement. Optogenetic modulation of layer 4/5 PV+ neurons in restrained animals will be required to provide supporting evidence for sharply tuned FS HD cells to participate in improved HD tuning in principal cells. Gain control is another proposed function for inhibition in the cortex. Bidirectional optogenetic modulation of PV+ neurons imposed gain control in visual and auditory systems instead of drastically changing principal cell tuning (Atallah et al., 2012; Moore and Wehr, 2013; Wilson et al., 2012). Sharp tuning of putative inhibitory cells described here may relate to the need to decrease the gain of HD signal within the S1HL, which is compatible with the supposition that spatial representation in the S1 may relate to body parts in space, rather than the whole organism (Brecht, 2017; Long and Zhang, 2018). Alternatively, it has been shown in the vibrissae S1, thalamic-mediated feedforward inhibition is key to suppress motor contributions to somatosensation (Yu et al., 2016). In this scheme, we assume motor inputs at least partially drive spatial responses in the S1HL; sharply tuned FS HD cells may provide the strong inhibition to delineate current HD (sensory) from future (motor) HD.

In this study, we carried out the first in-depth characterization of spatial representation in the S1HL (Long and Zhang, 2018). We show a relatively high proportion of putative inhibitory FS cells code for HD with better precision than their RS, putative principal cell counterparts. These data challenge the prevailing view of cortical FS cell function in general, and how HD information is utilized in the brain. The unequivocal anatomical classification of reported FS HD cells is crucial for unlocking an apparent novel cortical mode of operation and (spatial) feature selection.

## Materials and Methods

### Subjects

Seven male Long-Evans rats (2-4 months old, 250-450 grams at the time of the surgery) were used for this study. All animals were housed in groups of four prior to surgery and singly housed in transparent cages (35 cm x 45 cm x 45 cm, W x L x H) and maintained on a 12-hour reversed light-dark cycle (lights on at 9 p.m. and off at 9 a.m.) after surgery. Experiments were performed during the dark phase. Rats were maintained in a vivarium with controlled temperature (19-22°C), humidity (55-65%). and were kept at about 85-90% of free-feeding body weight. Food restriction was imposed 8-24 hours before each training and recording trial. Water was available *ad libitum*. All animal experiments were performed in accordance with the National Animal Welfare Act under a protocol approved by the Animal Care and Use Committees from both Army Medical University and Xinqiao Hospital.

### Surgery and tetrode placement

Rats were anesthetized with isoflurane for implant surgery. The local anaesthetic lidocaine was applied to the scalp before the incision was made. Microdrives loaded with four tetrodes were implanted to target the hindlimb region of the primary somatosensory cortex (anterior-posterior (AP): 0.2-2.2 mm posterior to bregma; medial-lateral (ML): 2.2-3.4 mm lateral to midline, dorsal-ventral (DV): 0.4/0.6-3 mm below the dura.), secured with dental cement with 8-10 anchor screws. One screw served as the ground electrode. Rats were given Temgesic as post-op analgesic. Tetrodes were assembled with four 17 µm Platinum/Iridium wires (#100167, California Fine Wire Company). Tetrodes had impedances between 150 and 300 kΩ at 1 kHz through electroplating (nanoZ; White Matter LLC).

### Training and data collection

Behavioral training, tetrode advancement, and data recording started a week after surgery. Rats were trained to forage in a 1 m x 1 m square box with a white cue card (297 mm x 210 mm) mounted on one side of the wall. Food pellets were scattered into the arena intermittently to encourage exploration.

Each recording session lasted between 20 and 40 min to facilitate full coverage of the testing arena. Tetrodes were advanced in steps of 25 or 50 µm daily until well-separated single units can be identified. Data were acquired by an Axona system (Axona Ltd., St. Albans, U.K.) at 48 kHz, band-passed between .8-6.7 kHz and a gain of x 5-18k. Spikes were digitized with 50 8-bit sample windows. Local field potentials were recorded from one of the electrodes with a low-pass filter (500 Hz).

### Spike sorting, cell classification and rate map

Spike sorting was manually performed offline with TINT (Axona Ltd, St. Albans, U.K.), and the clustering was primarily based on features of the spike waveform (peak-to-trough amplitude and spike width), together with additional autocorrelations and cross-correlations (Skaggs et al., 1996; Wilson and McNaughton, 1993). During our manual cluster cutting, we always counted neurons with similar or identical waveform shapes only once across consecutive recording sessions. Putative fast-spiking interneurons were classified as cells with an average firing rate above 3 Hz and a peak-to-trough spike width below 300 µs (Peyrache et al., 2012).

Two small light-emitting diodes (LEDs) were mounted to the headstage to track the rats’ position and orientation via an overhead video camera. Only spikes with instantaneous running speeds > 2.5 cm/s were chosen for further analysis in order to exclude confounding behaviors such as immobility, grooming, and rearing (Zhang et al., 2013).

To classify firing fields and firing rate distributions, the position data were divided into 2.5-cm × 2.5-cm bins, and the path was smoothed with a 21-sample boxcar window filter (400 ms; 10 samples on each side) (Boccara et al., 2010; Zhang et al., 2013). Cells with > 100 spikes per session and with a coverage of >80% were included for further analyses. Maps for spike numbers and spike times were smoothed with a quasi-Gaussian kernel over the neighboring 5 × 5 bins (Zhang et al., 2013). Spatial firing rates were calculated by dividing the smoothed map of spike numbers with spike times. The peak firing rate was defined as the highest rate in the corresponding bin in the spatial firing rate map. Mean firing rates were calculated from the whole session data. Spatial autocorrelation was calculated with smoothed rate maps (Boccara et al., 2010; Zhang et al., 2013). Autocorrelograms were derived from Pearson’s product-moment correlation coefficient correcting for edge effects and unvisited locations.

### Analysis of head direction cells

The rats’ head direction was estimated by the relative position of the LEDs differentiated through their sizes (Taube, 1995; Zhang et al., 2013). The directional tuning curve for each recorded cell was drawn by plotting the firing rate as a function of the rat’s head angle, which is divided into bins of 3 degrees and then smoothed with a 14.5-degree mean window filter (2 bins on each side). To avoid bias, data were only used if all head angle bins contain data (Zhang et al., 2013).

The strength of directionality was calculated by computing the mean vector length from circular distributed firing rates. The chance values were determined by a shuffling process, with the entire sequence of spike trains time-shifted between 20 s and the whole trail length minus 20 s along the animals’ trajectory. This shuffling process was repeated 100 times for each cell, generating a total of 210,700 permutations for the 2107 somatosensory neurons. This shuffling procedure preserved the temporal firing characteristics in the unshuffled data while disrupting the spatial structure at the same time. Cells were defined as head direction cells if the mean vector lengths of the recorded cells were larger than the 99^th^ percentile of the mean vector lengths in the shuffled distribution.

### Analysis of angular head velocity cells

The modulation of firing rate by animals’ angular head velocity (AHV) was calculated as previously described (Taube, 1995). Briefly, the first derivative of head direction (angular velocity) for each time sample was computed. For each cell, the firing rate was plotted as a function of AHV in 6 degree/s bin. To minimize the sampling bias, bins with less than 50 samples per second were excluded. The AHV score was defined by calculating the Pearson’s correlation coefficient between the angular head velocity and firing rate. Shuffling was performed in the same procedure used for defining head direction cells. Cells were defined as AHV cells if the AHV scores of the recorded cells were larger than 0.5.

### Analysis of theta modulation and power spectral density

To calculate fluctuations of neural activity through the theta cycle, we offline-filtered local field potentials (LFPs) to extract S1HL theta oscillations. For the low-pass filtering, 4 and 5 Hz were selected as stopband and passband low cut-off frequencies, respectively, while 10 and 11 Hz were selected as passband and stopband high cut-off frequencies, respectively. Theta modulation was calculated from the fast Fourier transform (FFT)-based power spectrum of the spike-train autocorrelation. When the mean spectral power within 1 Hz range of the theta peak within the 4-11 Hz frequency range was at least 5 times larger than the mean spectral power from 0 Hz to 125 Hz, the cell was classified as theta modulated. To quantify the theta rhythmicity of the local field potentials (LFP) within the somatosensory cortex, the LFP was bandpass-filtered with the theta range (4-11 Hz) to acquire theta-filtered LFP traces. Power spectral density was computed with FFT.

### Analysis of Bursty Firing Properties

We analyzed the interspike interval (ISI) histogram to determine the bursty discharge patterns as described previously (Ebbesen et al., 2016). ISI probability distribution was first computed for each head direction cell by binning the ISIs below 60 ms with 1 ms bins and normalized that the area under the curve equals 1. Then a principal component analysis (MATLAB, MathWorks) was performed on the ISI probability distributions for all the neurons and the first two principal components (PC1 and PC2) were obtained, followed by a 2D Gaussian kernel smoothed density estimate (Botev et al., 2010).

### Cross-correlogram and putative synaptic connections

We identified putative monosynaptic connections by using spike-time cross-correlograms as described by others (Bartho et al., 2004; Csicsvari et al., 1998). Briefly, short-latency (< 4ms) peaks with the amplitude above 5 SDs of baseline mean of the cross-correlograms were considered as putative monosynaptic excitatory connections. Similarly, short-latency (< 4ms) troughs with the amplitude below 5 SDs of baseline mean of the cross-correlograms were considered as putative monosynaptic inhibitory connections. For simultaneously recorded neuron pairs on the same tetrode, the 0-1 ms bins of the cross-correlogram were not considered due to the unresolved superimposed spikes.

### Histology and reconstruction of recording positions

At the end of the experiment, rats were euthanized with an overdose of sodium pentobarbital and perfused transcardially with phosphate-buffered saline (PBS) followed by 4% paraformaldehyde (PFA). Brains were removed and stored in 4% PFA overnight. The brain was then placed in 10, 20 and 30% sucrose/PFA solution sequentially across 72 hours before sectioning using a cyrotome. Thirty-micron sections were obtained through the implant region. Sections were mounted on glass slides and stained with cresyl violet (Sigma-Aldrich). The final recording positions were determined from digitized images of the Nissl-stained sections. Positions of each recording were estimated from the deepest tetrode track, notes on tetrode advancement with tissue shrinkage correction by dividing the distance between the brain surface and electrode tips by the last advanced depth of the recording electrodes. All electrode traces were confirmed to be located within the S1defined by The Rat Brain Atlas (Paxinos and Watson, 2007).

## Acknowledgments

We are indebted to H. Wu, W. Tang, Q. Ren, H. Chen and S. Lv for their encouragement and generous help. We thank J. Cai and B. Deng for their excellent technical assistance. X.L. is supported by the Chongqing Municipality postdoctoral fellowship (Grant# cstc2019jcyj-bshX0035). This work is supported by the National Natural Science Foundation of China through the Project Grant NSFC-31872775 and startup funds from Xinqiao Hospital (Grant# 2017A034 and Grant# 2019XQY16) and Army Medical University (Grant# 2017R028) to S.-J.Z.

## Author Contributions

S.-J.Z. conceived the project. X.L., C.K.Y. and S.-J.Z. designed the study. X.L. and S.-J.Z. performed the experiments and collected the data. X.L. and C.K.Y. analyzed the data and made the figures. X.L., C.K.Y. and S.-J.Z. wrote the manuscript.

## Competing interests

The authors declare no competing interests.

## Supplementary Information

This file contains Supplementary figures S1-S8.

## Data and Code Availability

The data and source code that support the findings of this study are available from the corresponding author upon reasonable request.

---

**Supplementary Fig. S1:**
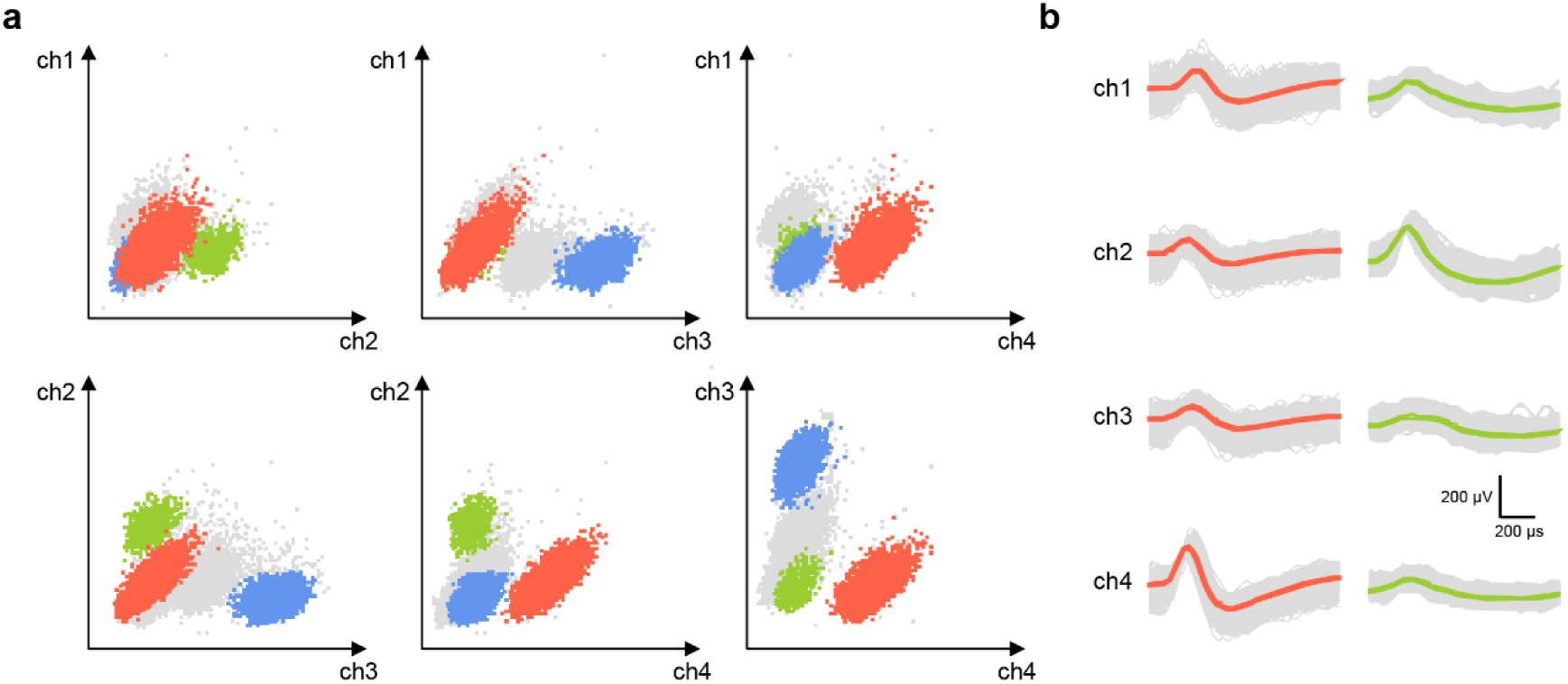
Cluster diagrams and waveforms recorded from the hindlimb region of the primary somatosensory cortex. (**a**) Scatterplots show the relationship between peak-to-trough amplitudes for all spikes, which are recorded on six types of permutation and combination from all of four tethered electrodes (ch1-ch4) on a specific tetrode. Each dot represents a single recorded spike. (**b**) Waveforms from two separated FS (red) and RS (green) clusters in the scatterplots are shown from four electrodes on the same recorded tetrode based on the shape of waveforms and the average firing rates. The cluster diagrams show spike clusters are well separated from our recordings in the S1HL.

**Supplementary Fig. S2:**
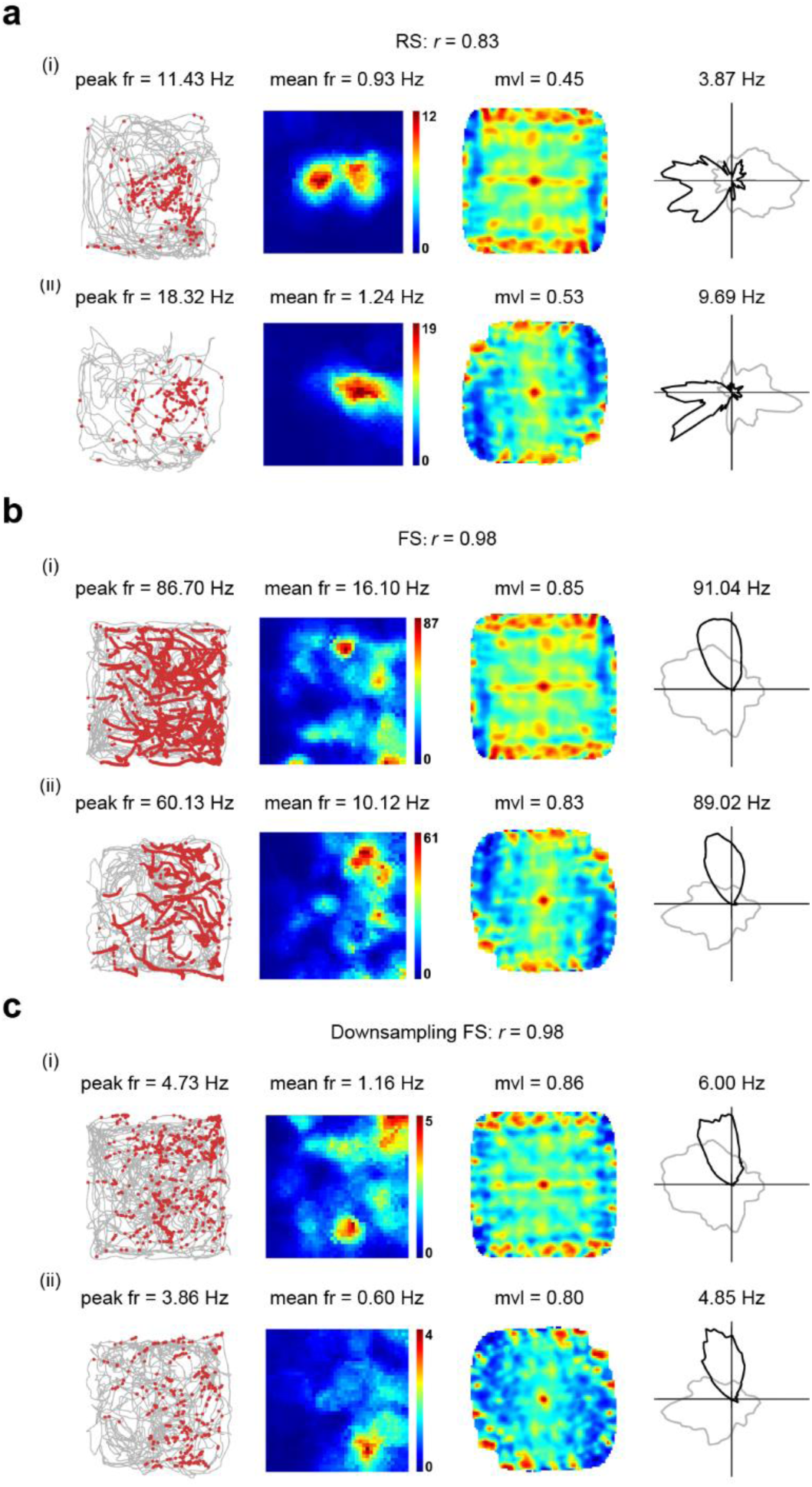
Angular stability of S1HL head direction cells. **a**-**c**, Intra-trial angular stability between the first and second halves of RS, FS and downsampled FS HD cells from **Fig. 2a**. From left to right: trajectory (grey line) with superimposed spike locations (red dots); smoothed rate maps; spatial autocorrelation; and HD tuning curve (black) plotted against dwell-time (grey) for the first half (**i**) and the second half (**ii**) of the trials. Firing rate is color-coded with blue indicating minimum firing rate and red indicating maximum firing rate. The scale of the autocorrelation maps is twice that of the spatial firing rate maps. Peak firing rate (fr), mean fr, mean vector length (mvl) and angular peak rate for each representative HD cell are labeled at the top of the panels. Correlation coefficients of the distributed firing rate across all directional bins between the first and second halves of individual recording trials are indicated with *r*.

**Supplementary Fig. S3:**
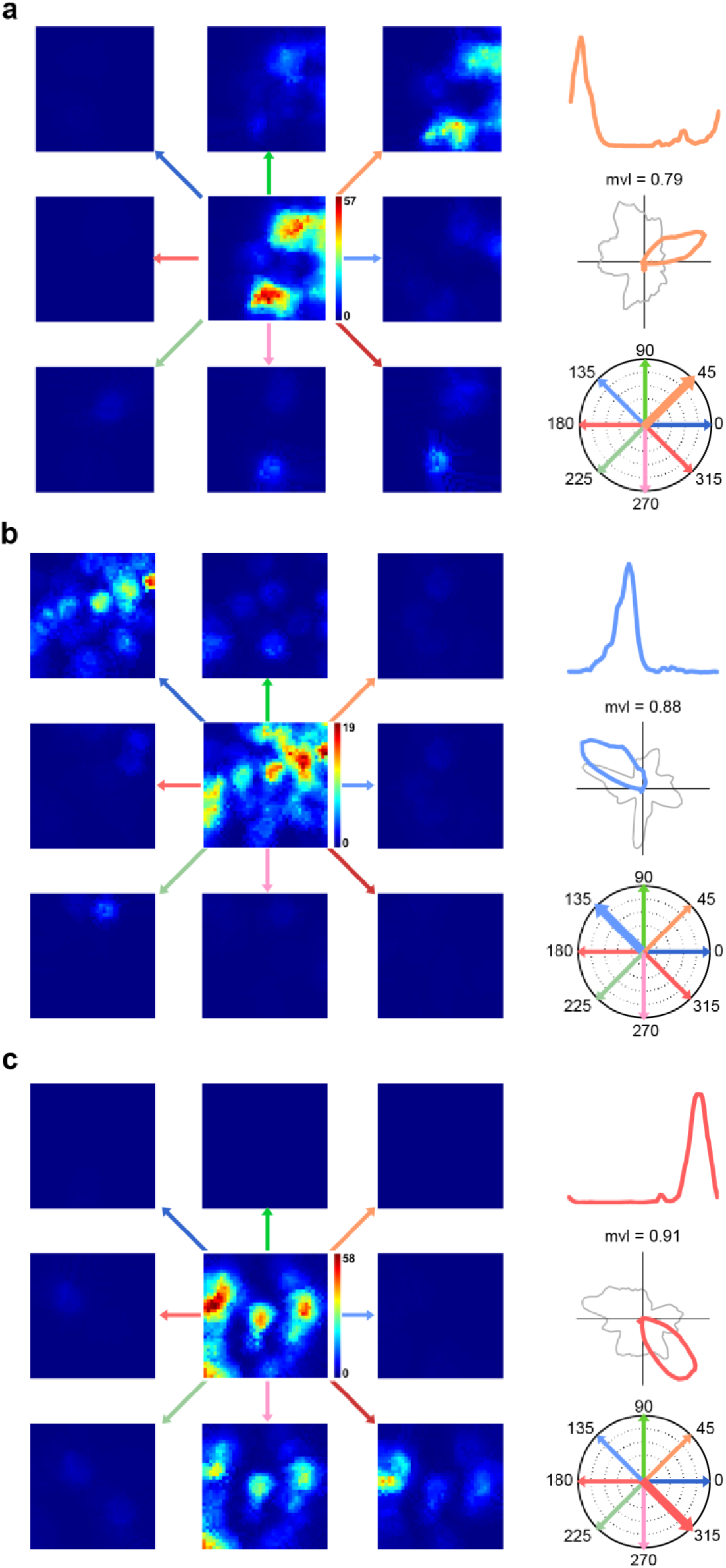
No positional bias in head direction tuning in S1HL FS head direction cells. **a-c**, Three examples of S1HL FS HD cells recorded from freely moving rats. For each recorded FS HD cell, 8 smoothed rate maps across all eight HD sectors are shown in the following plots. Left, one spatial firing rate map of a single FS HD cell is divided into eight smoothed rate maps across all eight HD sectors segmented by 45° each. Top right, HD tuning curve. Middle right, firing rate polar plots plotted against dwell-time polar plot (grey). Bottom right, circular distribution of the preferred firing direction across 360°.

**Supplementary Fig. S4:**
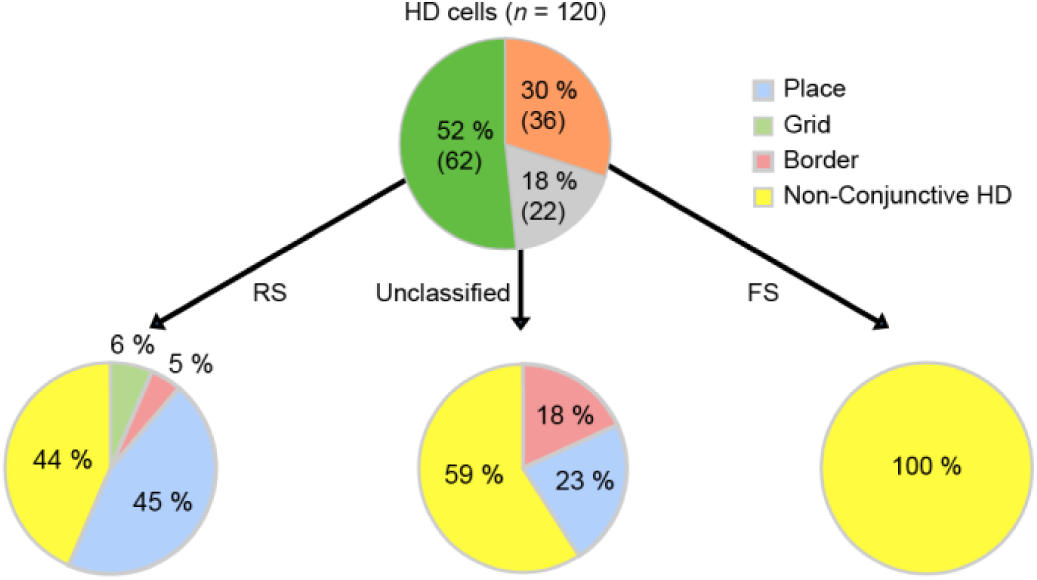
Classification of somatosensory head direction cells. Pie diagram showing the classification of fast-spiking (FS), regular-spiking (RS) and unclassified head direction (HD) cells into conjunctive and non-conjunctive HD cells.

**Supplementary Fig. S5:**
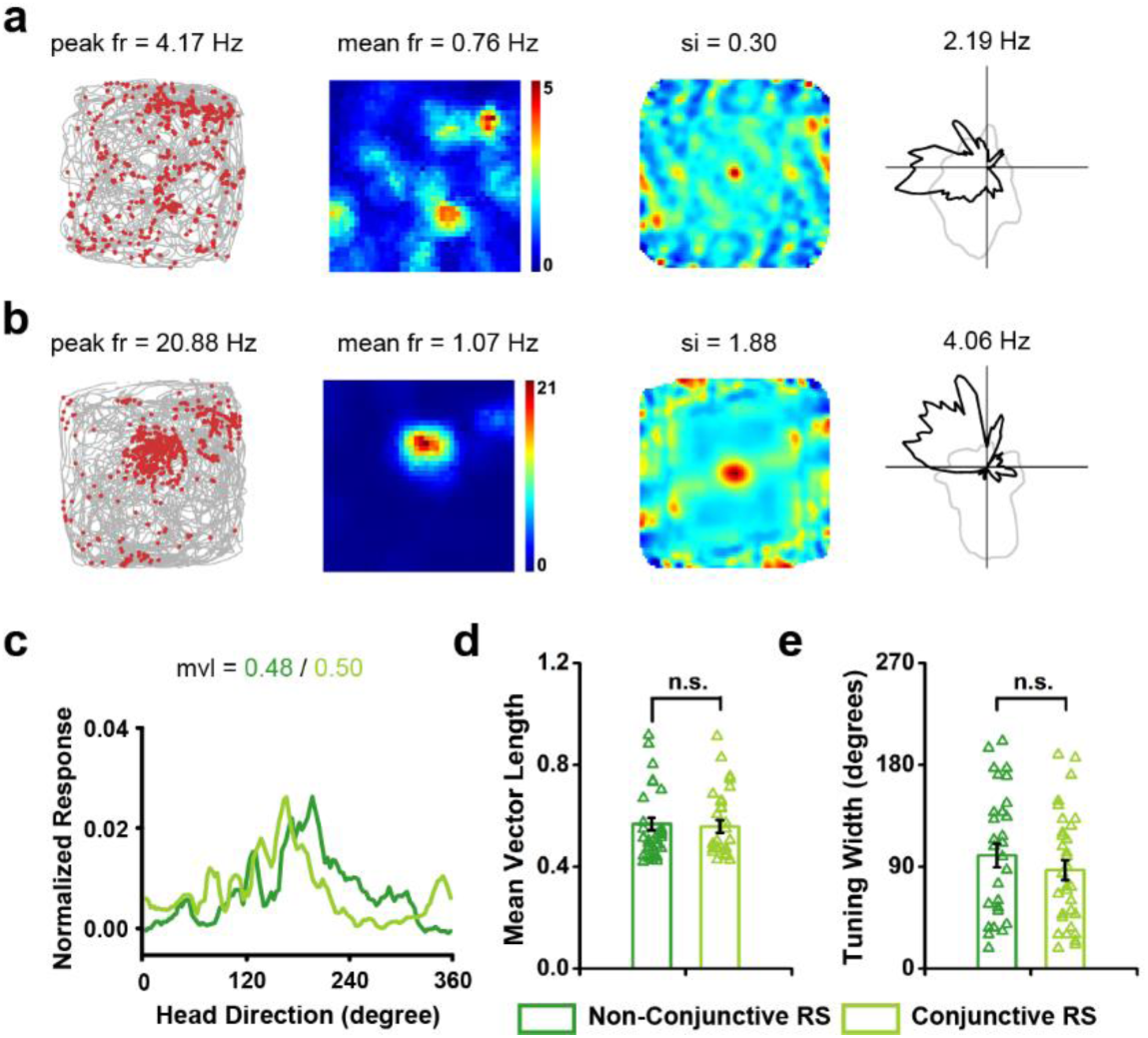
Conjunctive and non-conjunctive regular-spiking cells show similar head directional tuning. **a**, Representative non-conjunctive RS HD and **b**, conjunctive RS HD x place cells. From left to right: trajectory (grey line) with superimposed spike locations (red dots); smoothed rate maps; spatial autocorrelation; and HD tuning curve (black) against dwell-time (grey). **c**, Normalized head direction tuning curve for the non-conjunctive (dark green) and conjunctive (light green) RS HD cells. **d**, Non-conjunctive RS HD cells (dark green, *n* = 30) and conjunctive RS HD cells (dark green, *n* = 30) have an indistinguishable mean vector length. **e**, Non-conjunctive RS HD cells show comparable tuning width as conjunctive RS HD cells.

**Supplementary Fig. S6:**
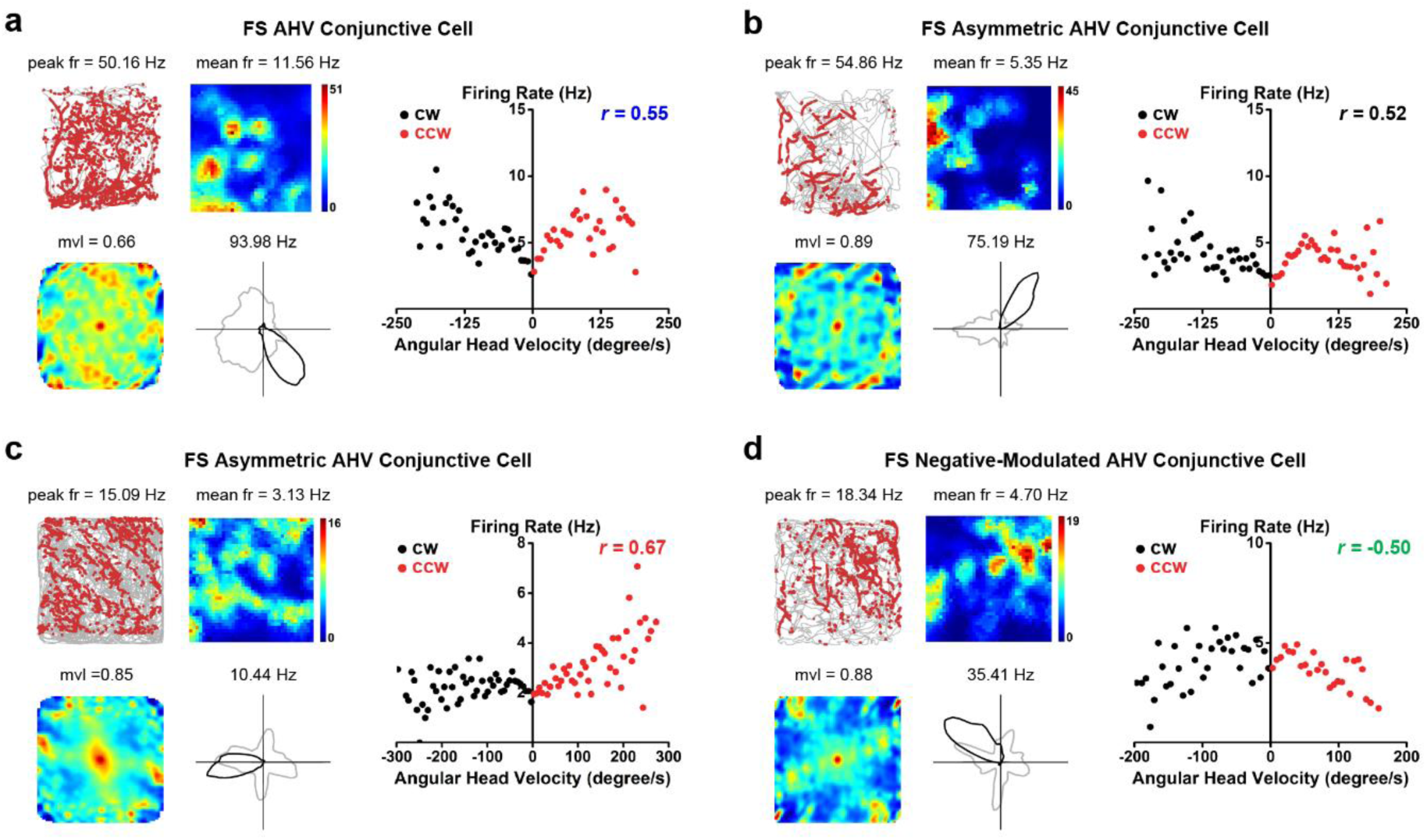
Conjunctive FS HD × AHV cells. **a-d**, Four representatives of conjunctive FS HD × AHV cells. **a**, FS HD cell with positive AHV modulation; **b**, FS HD cell with asymmetrical (CW) AHV modulation; **c**, FS HD cell with asymmetrical (CCW) AHV modulation; **d**, FS HD cell with symmetrical negative AHV modulation. Left panel: Trajectory (grey line) with superimposed spike locations (red dots; top left), smoothed rate maps (top right), spatial autocorrelation (bottom left) and HD tuning curve (black) plotted against dwell-time (grey; bottom right). Firing rate is color-coded with dark blue indicating minimal firing rate and dark red indicating maximal firing rate. The scale of the autocorrelation maps is twice that of the spatial firing rate maps. Peak firing rate (fr), mean fr, mean vector length (mvl) and angular peak rate for each representative head direction cell are labelled at the top of the panels. Right panel: Scatterplot of the firing rate versus the angular velocity of the representative cells.

**Supplementary Fig. S7:**
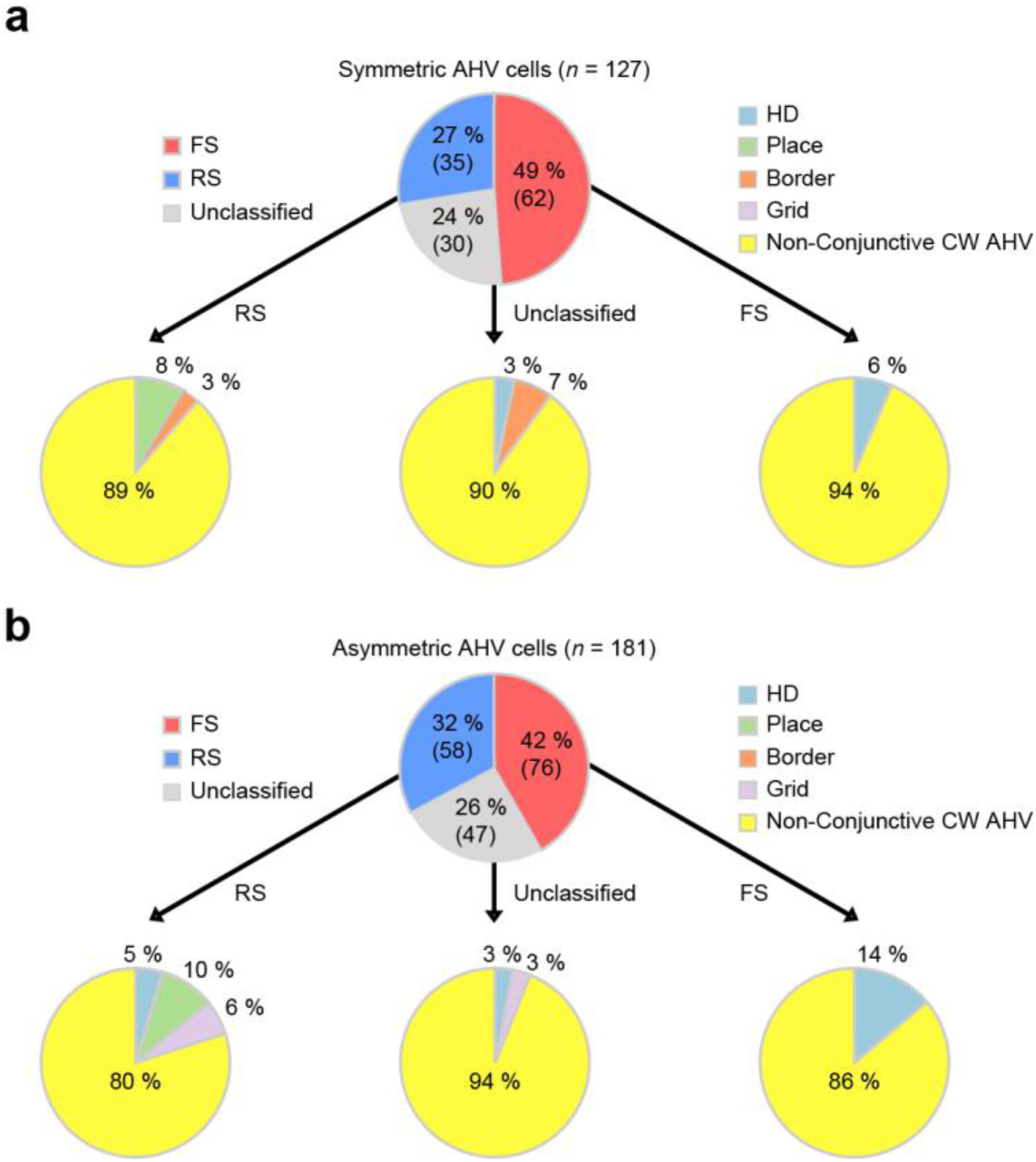
Classification of S1HL AHV cells. **a**, Pie diagram showing the composition of three neuronal types of symmetric AHV cells. **b**, Pie diagram showing the composition of three neuronal types of asymmetric AHV cells.

**Supplementary Fig. S8:**
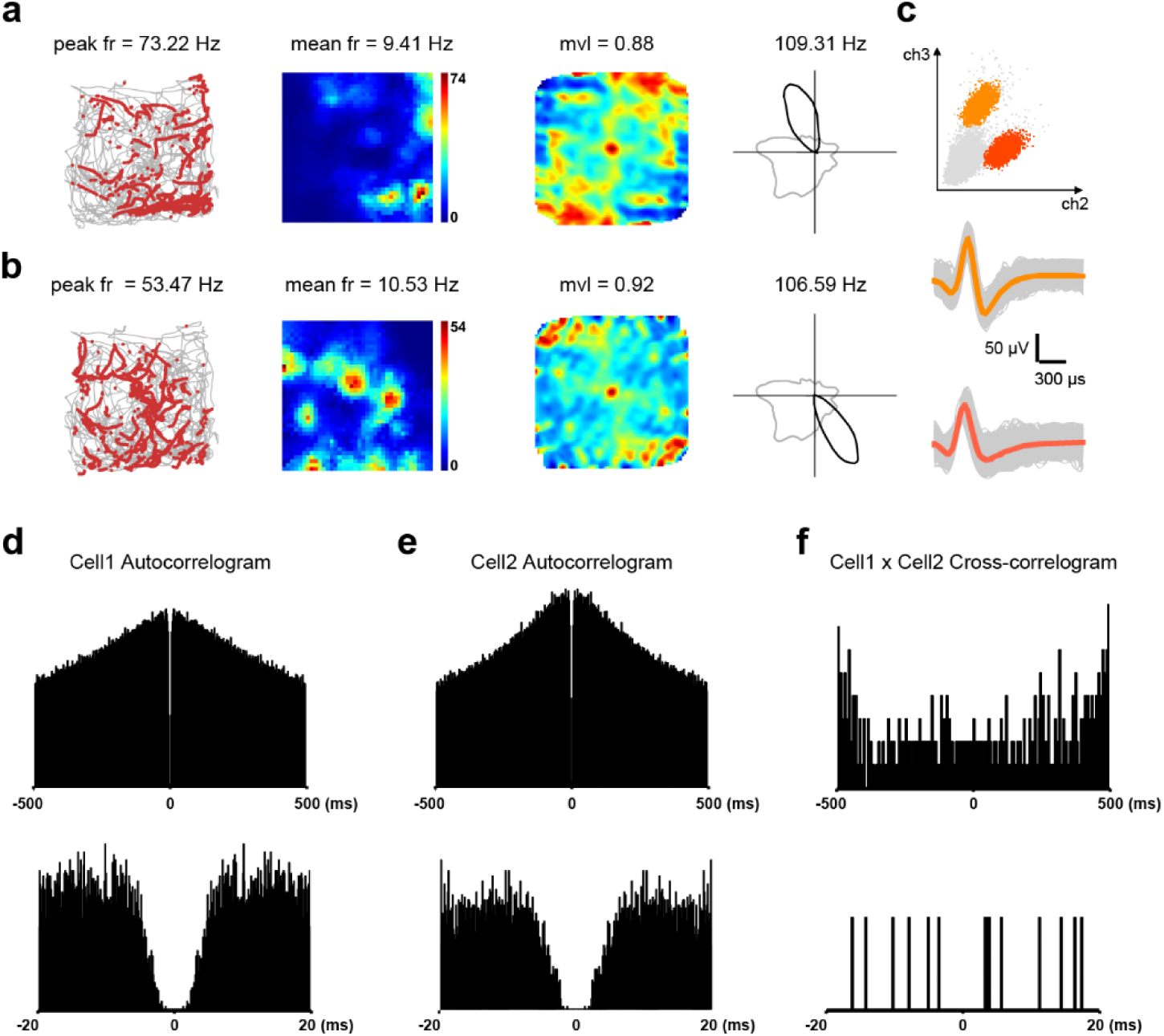
Two simultaneously recorded FS HD cells on the same tetrode. **a, b**, Two simultaneously recorded FS somatosensory HD cells from the same tetrode. Trajectory (grey line) with superimposed spike locations (red dots) (left column); spatial firing rate maps (middle left column), autocorrelation diagrams (middle right column), head direction tuning curves (black) plotted against dwell-time polar plot (grey) (right column). **c**, Cluster diagrams and waveforms. **d, e**, Representative 1000 ms spike-time autocorrelogram plot of the two representative somatosensory FS HD cells. **f**, Representative 1000 ms spike-time cross-correlogram of two exampled cells above. Bottom panels of **d-f**, shorter time scale of autocorrelogram and cross-correlogram.

